# Mitochondrial ribosome content as a proxy for respiration

**DOI:** 10.64898/2026.01.20.700566

**Authors:** Francis Mairet

## Abstract

Quantifying cellular activities remains a major challenge across fields ranging from microbial ecology to biotechnology and biomedical sciences. Building on the well-established linear relationship between growth rate and ribosome content — the so-called microbial growth law — this study proposes using organelle ribosome content to infer metabolic activity. In exponentially growing yeast, including under overflow conditions, a strong linear correlation was observed between mitochondrial ribosome content and oxygen uptake rate, demonstrating the robustness and potential of this approach. In addition, cytoplasmic and mitochondrial ribosome fractions exhibited a linear relationship, while overflow conditions appeared as outliers, providing a means to identify such metabolic states. Altogether, these findings highlight the promise of organelle ribosome quantification as a novel proxy for deciphering cellular metabolism.

## Introduction

Microorganisms are virtually ubiquitous. They play key roles in natural ecosystems and in the health of their hosts, as well as in numerous biotechnological processes. Over the past decade, interest in characterizing microbial communities has grown, for example through metabarcoding or metatranscriptomic approaches. However, quantifying microbial activity remains challenging, especially in natural environments. While progress has been made in mapping the spatial and temporal distribution of microorganisms, new approaches are still needed to uncover their functions and quantify the fluxes that drive ecosystem processes (Strzepek et al., 2022).

One approach proposed to estimate microbial growth rate relies on the so-called microbial growth law, a linear relationship between growth rate and ribosome content observed experimentally in bacteria and protists (Scott et al., 2010). This empirical relationship can be interpreted through the lens of resource allocation theory: assuming that microorganisms have evolved strategies that maximize their fitness, metabolism can be modeled as an optimization problem in which resources (e.g., proteins) are allocated among different functional sectors — substrate uptake, ATP production, protein synthesis, etc. — to maximize growth rate (Molenaar et al., 2009). This framework explains the microbial growth law: under nutrient limitation, more resources are devoted to transporters and catabolic enzymes, reducing those available to ribosomes. As nutrient availability increases, enabling faster growth, more resources are allocated to ribosome production (Scott et al., 2014; Bosdriesz et al., 2015). The approach has since been extended to include, for instance, the effects of temperature (Mairet et al., 2021) or antibiotics (Scott et al., 2014).

Building on this principle, ribosome content (or proxies such as total RNA or ribosomal protein expression) is now widely used to estimate in situ microbial growth rates and to gain insight into ecosystem functioning (Blazewicz et al., 2013). Although this approach requires careful interpretation, it provides a simple yet powerful tool to assess microbial growth in complex systems.

This raises an intriguing question: if ribosome abundance can serve as a reliable proxy for cellular growth, could the same logic be applied at the organelle level to infer specific metabolic activities? Beyond growth rate, quantifying metabolic fluxes remains a central challenge across fields from microbial ecology to biotechnology and biomedical sciences. A particularly relevant case is respiration. Under conditions of high substrate availability, some cells can achieve faster growth by repressing respiration and enhancing glycolysis—a phenomenon known as overflow metabolism. Observed in yeast (the Crabtree effect) and cancer cells (the Warburg effect), this behavior can be explained by cellular resource allocation (Molenaar et al., 2009). Here, we show that mitochondrial ribosome content correlates with oxygen uptake rate, providing a means to estimate respiration and to detect overflow metabolism. This observation opens new avenues for deciphering cellular activities through organelle ribosome content.

## Results

Experimental data from *Saccharomyces cerevisiae* (Elsemman et al., 2022) were analyzed, including cells grown either in batch culture with different carbon sources or in continuous culture under various levels of glucose limitation (see Method). Across all these conditions, a clear linear correlation was observed between mitochondrial ribosome content and oxygen uptake rate (Figure 1, R^2^ = 0.97). Interestingly, data points corresponding to overflow metabolism—characterized by high specific growth rate but low oxygen consumption—fall along the same trend, highlighting the consistency of this relationship regardless of the underlying metabolic regime.

**Figure 1.**
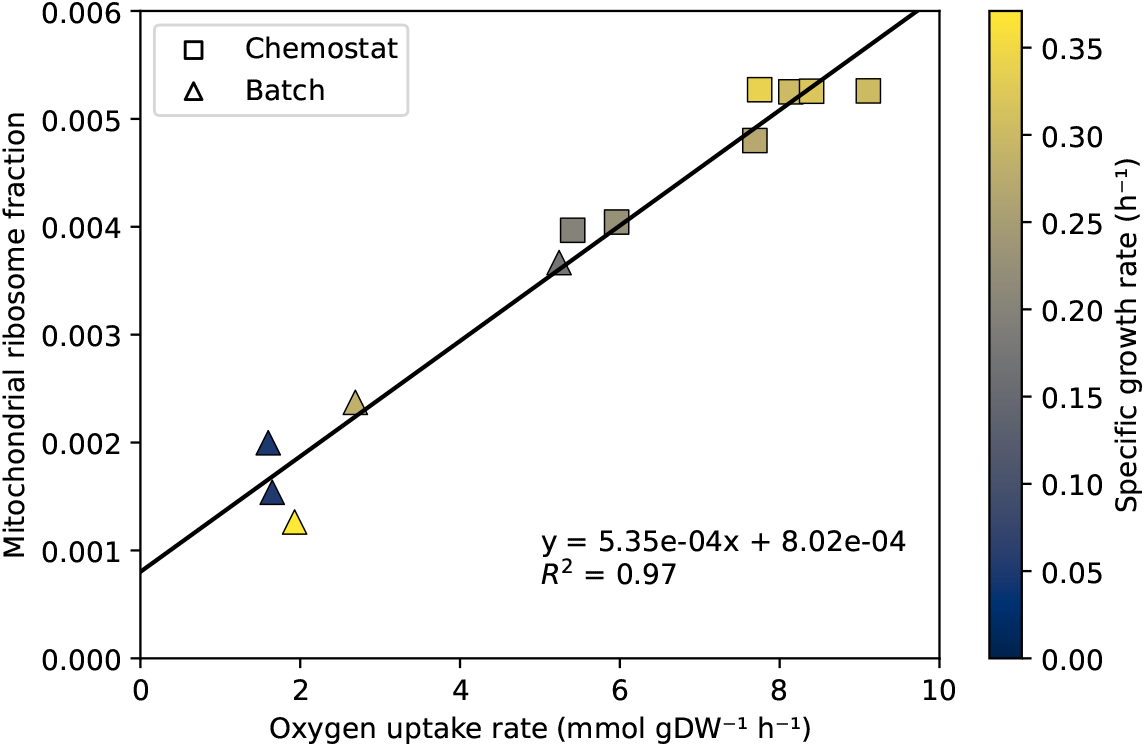
Correlation between the fraction of mitochondrial ribosomes and oxygen uptake rate in exponentially growing *Saccharomyces cerevisiae* under different carbon sources in batch culture (triangles) and varying dilution rates in glucose-limited chemostat (squares) (Elsemman et al., 2022). Colors indicate specific growth rates.

A few simple hypotheses are sufficient to explain this correlation. First, we assume that respiration is driven by the pool of mitochondrial-encoded enzymes and that, together with a subset of nuclear-encoded enzymes that are coordinately expressed, these respiratory enzymes operate near saturation (Bennett et al., 2009). At the coarse-grained level, the oxygen uptake rate *q*_*O*2_ is therefore proportional to the fraction of mitochondrial-encoded enzymes *m*:

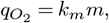

where *k*_*m*_ is the maximum rate constant of respiration. The fraction of mitochondrial enzymes *m* results from a balance between synthesis (at a rate *ρ*_*m*_) and degradation (at a first-order rate *λ*), assuming that dilution due to growth is negligible:

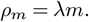

The synthesis rate *ρ*_*m*_ is driven by mitochondrial ribosomes, representing a fraction of the proteome *r*_*m*_. These ribosomes translate at a constant rate *k*_*r*_, near saturation, so the synthesis rate of mitochondrial enzymes depends only on the mitochondrial ribosome fraction:

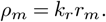

Combining these three equations yields a direct linear relationship between mitochondrial ribosome content and oxygen uptake rate:

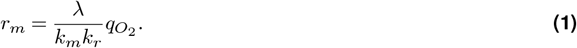

A rough estimate of the parameter 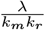 based on literature values gives 1.40 g.h.mol^−1^ (see Method), in reasonable agreement with the slope of the experimental correlation (0.53 g.h.mol^−1^). The non-zero intercept indicates the existence of a minimal content of mitochondrial ribosomes, not predicted by this simplified reasoning. It could be explained, for instance, by a reserve of ribosomes maintained in anticipation of nutrient shifts—analogous to the offset observed between growth rate and cytoplasmic ribosome content (Mori et al., 2017). Alternatively, this intercept may reflect the requirement for mitochondria to sustain additional, non-respiratory metabolic functions, such as the synthesis of amino acids or hemes (Malina et al., 2018).

Having established the correlation between mitochondrial ribosomes and respiration, we next examined how this relationship aligns with cytoplasmic ribosomes. Using the same dataset, we searched for a pattern using outlier detection procedure. We observed a linear correlation between mitochondrial and cytoplasmic ribosome contents after excluding two outliers (Figure 2, R^2^ = 0.81). These two automatically detected points correspond to the highest oxygen yields (*>* 100 gDW/mol O_2_) in the dataset, characteristic of overflow metabolism, whereas all other poins exhibit oxygen yield below 50 gDW/mol O_2_.

**Figure 2.**
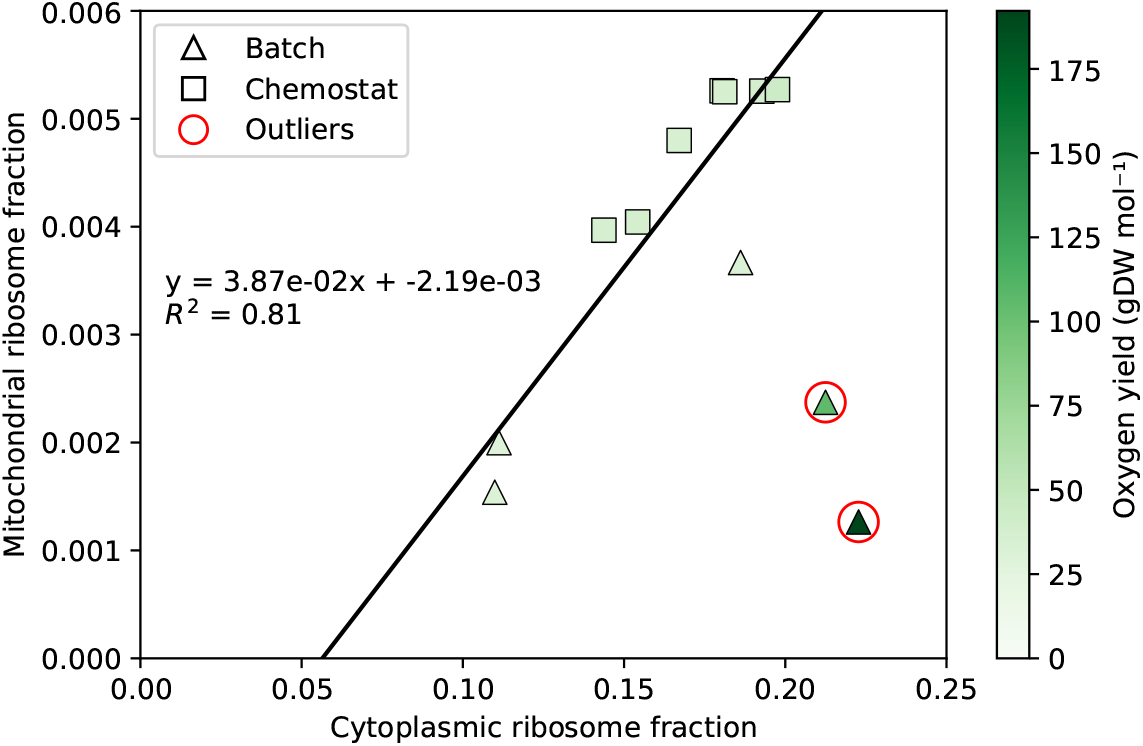
Correlation between mitochondrial and cytoplasmic ribosome fractions in exponentially growing *Saccharomyces cerevisiae* under different carbon sources in batch culture (triangles) and varying dilution rates in glucose-limited chemostat (squares) (Elsemman et al., 2022). Colors indicate oxygen yield. The automatically identified outliers, circled in red, correspond to overflow metabolism (i.e., high oxygen yield).

From the theoretical perspective, cytoplasmic ribosome content *r*_*c*_ is known to correlate with the specific growth rate *µ* (Scott et al., 2010):

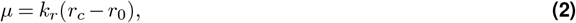

where *r*_0_ represents the minimal cytoplasmic ribosome content (Mori et al., 2017), and *k*_*r*_ is assumed here to be the same translation rate constant as for mitochondrial ribosomes, for simplicity.

Defining the oxygen yield coefficient as 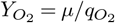, we obtain from Eq. (1) and Eq. (2):

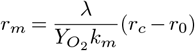

This expression captures the linear relationship observed between mitochondrial and cytoplasmic ribosome contents, assuming a constant oxygen yield. Data points falling below the regression line correspond to higher oxygen yield, indicative of overflow metabolism. All these theoretical predictions are consistent with the experimental data (see Figure 2). Additionally, the predicted slope of 5.24 *×* 10^−2^, obtained using literature parameter values (see Method) and the mean oxygen yield coefficient of the inlier points, agrees well with the experimentally determined slope of 3.87 *×* 10^−2^.

## Discussion

In the same way that cytoplasmic ribosome content correlates with specific growth rate, we have shown here a linear relationship between mitochondrial ribosome content and oxygen uptake rate in yeast. We therefore propose that mitochondrial ribosome abundance can serve as a proxy for respiratory activity. Our results also demonstrate that this approach can detect overflow metabolism by identifying outliers that fall below the linear correlation between cytoplasmic and mitochondrial ribosome contents established under non-overflow metabolic conditions.

These relationships can be explained through a few simple theoretical assumptions on cellular growth. A central hypothesis is that both respiratory enzymes and ribosomes operate near saturation, as observed for most enzymes in exponentially growing *Escherichia coli* (Bennett et al., 2009). This behavior likely reflects cellular economics (Molenaar et al., 2009; Bosdriesz et al., 2015): cells adjust their proteome composition to achieve an optimal allocation of resources, both at a coarse-grained level—between major cellular functions—and at a finer level, within each function, among individual enzymes. This optimization maintains a balanced ratio between metabolites and enzymes to maximize growth, thereby leading to near-saturation operation of enzymes and ribosomes.

In our theoretical framework, several processes are neglected, such as enzyme dilution due to growth (considered negligible relative to enzyme degradation) and non-respiratory oxygen consumption (e.g. in the synthesis of sterols or unsaturated fatty acids (Rosenfeld and Beauvoit, 2003)). These simplifying assumptions yield a tractable model that nonetheless captures the essential link between mitochondrial ribosome content and respiration. A main advantage of focusing on ribosome content, rather than on any specific protein, is that it provides a more integrative measure, further benefiting from the fact that translation rates are relatively conserved, whereas enzyme catalytic rates vary over several orders of magnitude (Milo and Phillips, 2015).

Future work should aim to validate these linear correlations across a broader range of species and experimental conditions. The increasing availability of in situ data—particularly metatranscriptomic datasets—offers promising opportunities for applying this method in complex environments. However, datasets that simultaneously combine gene expression profiles with quantitative flux measurements remain scarce. Such integrated datasets are crucial for further validating our approach and connecting biomolecular patterns with resulting metabolic rates (Strzepek et al., 2022).

Beyond unicellular organisms, emerging evidence suggests that this concept may have a broader applicability. For example, in human cells, genes involved in mitochondrial translation are overexpressed in tissues with high respiratory activity, such as muscle and heart (Jiang et al., 2020). Exploring whether similar relationships apply to cancer cells—known for their reliance on overflow metabolism—would be particularly insightful, in line with the ongoing search for empirical growth laws in tumor metabolism (Kochanowski et al., 2021).

Finally, this framework could be further extended by considering ribosomes from another organelle, the chloroplast. Together, these perspectives open new avenues for using organelle ribosome content as a quantitative proxy to decipher cellular metabolic activities.

## Method

### Data set

Experimental data were obtained from Elsemman et al. (2022). Two types of cultures were analyzed: batch cultures with different carbon sources (trehalose, galactose, maltose or glucose), and glucose-limited chemostat cultures with varying dilution rates. For each condition, the fraction of mitochondrial or cytoplasmic ribosomes was calculated by dividing the sum of all mitochondrial or cytoplasmic ribosomal protein abundances by the sum of all proteins.

### Data analysis

Linear regressions were performed using Python 3.13, with the coefficient of determination (*R*^2^) used to evaluate the quality of the correlation. The RANSAC algorithm from Scikit-Learn 1.7.2 (Pedregosa et al., 2011) was applied to automatically detect outliers.

### Parameter values

The mean degradation rate for mitochondrial proteins was taken from Christiano et al. (2014):

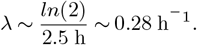

For ribosomes, we assume a translation rate of 10 amino acids per second, with average amino-acid and ribosome masses of 100 g.mol^−1^ and 2.7 *×* 10^6^ g.mol^−1^, respectively (Milo and Phillips, 2015). The translation rate, expressed as the mass of protein synthesized per mass of ribosomal protein per unit time, is given by:

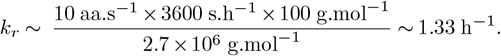

For the respiration rate, considering the eight enzymes encoded by the mitochondrial genome (Malina et al., 2018) with a total mass of ~ 244 kDa and a mean catalytic rate of 10 s^−1^ (Milo and Phillips, 2015), we obtain:

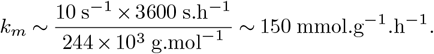

## Data, Materials, and Software Availability

Experimental data were taken from Elsemman et al. (2022). The script for data analysis and figure plotting is available on GitHub: https://github.com/fmairet/mitoribosome.

## Acknowledgments

I thank all my colleagues for the helpful discussions that supported this work.

## Notes

https://github.com/fmairet/mitoribosome

### Competing Interest Statement

The authors have declared no competing interest.

